# Electrophysiological correlates of divergent projections in the avian superior olivary nucleus

**DOI:** 10.1101/2024.03.06.583566

**Authors:** James F. Baldassano, Katrina M. MacLeod

## Abstract

The physiological diversity of inhibitory neurons provides ample opportunity to influence a wide range of computational roles through their varied activity patterns, especially via feedback loops. In the avian auditory brainstem, inhibition originates primarily from the superior olivary nucleus (SON) and so it is critical to understand the intrinsic physiological properties and processing capabilities of these neurons. Neurons in the SON receive ascending input via the cochlear nuclei: directly from the intensity-coding cochlear nucleus angularis (NA) and indirectly via the interaural timing nucleus laminaris (NL), which itself receives input from cochlear nucleus magnocellularis. Two distinct populations of SON neurons provide either inhibitory feedback to ipsilateral NA, NL, and the timing cochlear nucleus NM, or to the contralateral SON. To determine whether these populations correspond to distinct response types, we investigated their electrophysiology in brain stem slices using patch clamp electrophysiology. We identified three phenotypes: single-spiking, chattering tonic, and regular tonic neurons. The two tonic phenotypes displayed distinct firing patterns and different membrane properties. Additionally, fluctuating “noisy” currents were used to probe the capability of SON neurons to encode temporal features. Each of the three phenotypes had a distinct level of sensitivity to this temporally modulated input. By using cell fills and anatomical reconstructions, we could correlate the firing phenotypes with their axonal projection patterns. We found that SON axons exited via three fiber tracts with specific phenotypes comprising each tract. Two ipsilateral tracts exit the SON, with the laterally projecting tract composed entirely of regular tonic neurons and medially projecting tract composed of both regular and chattering tonic neurons. The contralateral projection was composed entirely of single spiking neurons. These results provide a basis for understanding the role of specific inhibitory cell types in auditory function and elucidate the organization of the SON outputs.

## Introduction

Inhibition is crucial for processing auditory information in central nervous system circuits^1–2,3,4^. In the auditory brainstem, inhibition sharpens phase locking for frequency tuning^5,6,7–10^, boosts gap detection capability^11,12^, and improves amplitude modulation coding^3,13^. The firing rates of inhibitory neurons in early auditory structures are driven by sound intensity^14–18^, thus their feedback can function as an intensity-dependent gain-control mechanism that regulates disparities in binaural sound levels (See reviews^19, 20^). Despite the multifaceted roles of inhibition, birds do not rely much on local inhibitory interneurons in their auditory brainstem nuclei. Instead, inhibition originates primarily in a nucleus, the superior olivary nucleus (SON)^21–24^, that is anatomically segregated from the ascending excitatory nuclei. To determine how the SON may underlie the multiple functions of inhibition in the auditory brainstem, we examined the intrinsic physiological properties of SON neurons.

In birds, the auditory nerve synapses onto two anatomically distinct cochlear nuclei: the intensity-coding nucleus angularis (NA) and the phase-, or time-, coding nucleus magnocellularis (NM)^25–27^. NM projects bilaterally to the nucleus laminaris (NL), which performs coincidence detection to calculate interaural time differences (ITD)^28^. Both NA and NL have excitatory projections to the ipsilateral SON^29,30^, which in turn projects either to the ipsilateral NM, NL, and NA, or to the contralateral SON (SONc). These projections come from two, non-overlapping populations of SON neurons^31^. *In vitro* studies of SON neurons have revealed two response types: 1) a single spiking phenotype wherein a single action potential is fired at the onset of a current injection, and 2) a tonic spiking phenotype wherein a series of action potentials are produced throughout the duration of the current injection^16,17,32^. It was unclear whether these two response phenotypes underlay the divergent projections from SON onto its targets.

*In vivo* recordings showed that SON neurons could generate a number of firing patterns in response to simple tone or noise sound stimuli. In these studies, most SON units were classified as either “onset” and produced action potentials only at the beginning of sound presentation, or “sustained” and produced action potentials throughout sound presentation^17^. These were primarily monaural responses, but some studies displayed subset had similar excitatory drive from both ipsilateral and contralateral stimuli^14^. While in earlier studies SON neurons were not thought to encode temporal features of sounds^5,16,23^, more recent work has demonstrated robust phase locking in sustained responses^17^. The underlying drivers of these variations could be from differential weighting of inputs from either NA or NL, variations in synaptic properties, or, as shown in both mammalian and avian cochlear nucleus neurons, diversity of intrinsic membrane properties^33–39^. In fact, temporally responsive intrinsic properties of a subset of NA neurons allow them to reliably encode features of the acoustic envelope^39–43^. We hypothesized that intrinsic properties of SON neurons were the primary driver of their temporal responsiveness. Furthermore, if we found that SON biophysical phenotypes corresponded to postsynaptic targeting, that might provide some functional insight into how diversity in SON could relate to the variety of roles for inhibition.

To investigate these properties *in vitro*, we used naturalistic current steps to expand the characterization of SON intrinsic phenotypes and axonal trajectory tracing to determine the cell-type specific projections of SON neurons. We demonstrate the presence of a second tonic firing phenotype that displays sensitivity to the temporal features of inputs. Furthermore, we determined that the firing phenotype was highly indicative of the neuron’s axonal projection. These results suggest that efferent and contralateral inhibitory signals may reflect different functional outputs and that intrinsic features of SON neurons may drive the different temporal properties seen *in vivo*.

## Materials and Methods

### Brain slice preparation

All animal procedures were performed with Institutional Animal Care and Use Committee approval and according to University of Maryland guidelines on animal welfare. White leghorn chicken (*Gallus gallus*) embryos (sex undetermined) incubated to embryonic day 17-18 were cooled and rapidly decapitated, and the head section containing the brain stem blocked, placed in a chilled, oxygenated low-sodium artificial cerebrospinal fluid (ACSF) (low-Na+ ACSF; in mM: 97.5 NaCl, 3 KCl, 2.5 MgCl_2_, 26 NaHCO_3_, 2 CaCl_2_, 1.25 NaH_2_PO_4_, 10 dextrose, 3 HEPES, and 230 sucrose) and dissected out of the cranium. The brain stem tissue block was mounted with cyanoacrylate glue and supported with 5% agarose gel. Standard transverse slices (250 μm thick) containing SON were cut on a vibrating tissue slicer (Leica Microsystems, Wetzler, Germany). For neuronal reconstruction experiments, “wedge” slices were cut to preserve the maximal amount of the recorded neuron’s axonal projection. After advancing through the tissue and just prior to collecting the section containing SON, the mounting platform was angled approximately 15°, such that the slice containing SON was approximately 400µm at the dorsal edge and 200µm at the ventral edge. We used wedge slices with a thick dorsal end and a thinner ventral end as opposed to slices that were uniform thickness in order to allow for maximum visibility under DIC within the SON while recording. Slices were incubated in normal ACSF (in mM: 130 NaCl, 3 KCl, 2 MgCl_2_, 26 NaHCO_3_, 2 CaCl_2_, 1.25 NaH_2_PO_4_, 10 dextrose, and 3 HEPES) at 34°C for 30 min then held at room temperature in normal ACSF until recording.

### Whole cell patch-clamp electrophysiology

Slices were placed in a recording chamber and continuously perfused with oxygenated, warmed normal ACSF (1–2 mL/min, 28-30℃) containing synaptic receptor blockers (3 µM strychnine, 20 µM SR95531 (gabazine), 15 µM 6,7-dinitroquinoxaline-2,3-dione (DNQX), and 20 µM D-AP5 (D-2-amino 5-phosphovalerate); Sigma) to isolate intrinsic activity. Whole cell patch-clamp recordings were performed on visually identified NA cells using infrared differential interference contrast video microscopy.

Initial glass recording pipette resistances were 3–7 MΩ. Pipettes were filled with a potassium gluconate intracellular recording solution (in mM: 110 potassium gluconate, 20 KCl, 1 EGTA, 2 MgCl_2_, 10 HEPES, 2 Na_2_ATP, 0.3 Na_2_GTP, 10 phosphocreatine and 0.2% biocytin). Series resistance, cell capacitance, and resting membrane voltage were measured upon break-in in voltage clamp mode. Voltage recordings were made with a MultiClamp 700B amplifier (Molecular Devices, Sunnyvale, CA) set to current-clamp mode. Application of the current stimulus and recording of the voltage output were controlled by an analog-to-digital board (National Instruments, Austin, TX) and a computer running custom software written in IGOR Pro (WaveMetrics, Lake Oswego, OR). A holding current was applied to maintain a constant voltage baseline of approximately −65 mV. Cell type classification protocols were carried out to identify neuronal phenotypes in SON with 400-ms-duration flat current steps of varying amplitudes from −150 pA up to 550 pA in 50 or 100 pA intervals. We collected data from 93 SON neurons (34 single spike, 23 regular tonic, 34 chattering tonic) across experiments.

### Regularity analysis

Tonic neurons fired multiple action potentials throughout the duration of a flat current injection but could be subdivided into at least 2 subtypes of tonic firing-regular and chattering based on the regularity of their spiking. Regularity was estimated from the coefficient of variation (CV) of the interspike intervals over the 400-ms response duration, such that the closer the CV was to zero, the more regular the firing. CV was calculated as the ratio of the standard deviation(σ) of the intervals over the mean (μ) of the intervals:

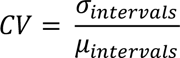

We used the CV for the first step above rheobase when determining the cell type, because that is the current level where the patterning emerges.

### Acquisition of f-I Curve and Analysis

Input-output functions were assessed by measuring firing-current (f-I) curves with standard flat current steps (400-ms duration as for classification protocols) or noisy current stimuli (2-s duration), as previously described^41,43^. Noise stimuli were constructed by convolving Gaussian white noise with an exponential function (time constant, 3-ms) added to a direct current (DC) step function. These noise stimuli simulate the arrival of many small, stochastic, and statistically independent synaptic currents, both excitatory and inhibitory. To accommodate cell-to-cell differences in input resistance, a standard noise current was created by calibrating the standard deviation (around a zero mean) of the noise current to generate a 2-mV standard deviation in the membrane potential in the target neuron, designated “1σ.” The amount of voltage fluctuation was varied by multiplying the standard noise stimulus by a factor of 2, 4, or 8 (i.e., 4-, 8-, and 16-mV voltage fluctuation, respectively). Each trial had an interstimulus interval of 8 seconds. A hyperpolarizing conditioning pre-pulse (−50 pA, 1-s duration) was applied before each stimulus to minimize sodium channel inactivation from the prior stimulus. A complete series of stimuli in the parameter space (mean DC, noise level) was systematically generated by varying the mean current amplitude and noise level independently (3 noise levels, 5–10 DC levels). Firing rates were averaged across one to three stimulus repetitions over the full 2 second duration and reported as means ± SD in Hertz.

Metrics to quantify differences between the lowest noise level (2σ) and highest noise level (8σ) curves were devised in Kreeger et al.^41^ and utilized here. One metric quantified the total area difference index (ADI) (28):

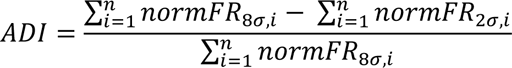

where normFR was the firing rate at each of n steps in each noise level normalized to the maximum firing rate in the whole data set. The second metric was the normalized difference in the maximum firing rate between 2σ and 8σ, ΔMaxFR:

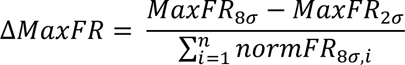

where MaxFR refers to the maximum firing rate value of each within-noise-level f-I curve. Both metrics varied from 0 (curves are identical) to 1 (maximum difference). Plotting these points (MaxFR index and the ADI index) against each other produced a spectrum of fluctuation sensitivity. Neurons closest to the origin were more “integrator-like,” whereas neurons further away from the origin were more “differentiator-like.” Using the criteria of ADI and ΔMaxFR > 0.2, we were able to test the noise f-I curves from 15 differentiators and 5 integrators with DTX. ΔMaxFR index and ADI index were statistically compared using Wilcoxon’s t test.

### Spike Timing Reliability

After noise was calibrated, a 4σ noisy current was injected into neurons at a single DC current step amplitude (typically 150–400 pA) to produce a firing rate between 20 and 50 Hz. Repeated trials of 2-s long frozen noisy current injections were injected (usually 30 to 45 trials, or until >600 spikes were collected) before and after DTX application. To quantify neuronal reliability, we calculated a shuffled autocorrelogram (SAC), a histogram of all interspike intervals between spikes across trials, excluding intervals within the same trial. The SAC is normalized by the normalizing factor (NF):

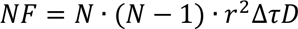

where N is the number of trials, r is the average firing rate, Δτ is the bin width of the correlation function (0.2-ms), and D is the length of the stimulus (2-s). The SAC was fit with a Gaussian function, whose central peak is defined as the correlation index (CI). The precision of the firing between trials is represented in the full width at half-maximum (FWHM) of the Gaussian curve, measured by

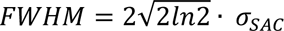

where σ_SAC_ represents 1 standard deviation of the Gaussian fit of the SAC. CI and FWHM, referred to as width, were quantified and compared using Wilcoxon’s t test. To determine the current signal that drove the response, the spike-triggered average (STA) was also calculated. A 60-ms snippet of the current stimulus, starting 50 ms preceding each spike, was extracted and averaged for all spikes within a set.

### Neuronal reconstruction and determination of postsynaptic target

Slices containing biocytin fills were fixed in 4% paraformaldehyde in 0.1 M phosphate buffer overnight and processed with Vectastain ABC (avidin-biotinylated HRP complex) Elite Kit (Vector Labs, Burlingame, CA). The shape of the slice, the boundary of SON, the location of filled neurons, and their morphological reconstructions were drawn using the computer-assisted tracing program Neurolucida (Microbrightfield, Williston, VT). Cytoarchitectural features and the shape of the slice were used to determine where the recording was in the rostral-caudal axis. Larger medial vestibular nuclei (MVN) and NM indicated more rostral and smaller, or a lack of, MVN and lower order nuclei (NA, NM, and NL) indicated more caudal.

Axons were able to be identified from dendrites due to their lack of branching, thinner diameter, and near constant diameter. Neurons were only used for analysis if their axon was a minimum of 50 micrometers in length and exited the boundary of the SON. Neuronal fills were superimposed onto a singular representative SON to determine emergent anatomical properties.

## Results

### SON contains heterogeneous electrophysiological phenotypes

Using simple current steps, we observed three action potential firing phenotypes in the voltage responses of SON neurons. First, a tonic firing phenotype (Fig. 1Ai) that fired tonically throughout the current step with firing rates that increased linearly with larger amplitude current steps (n=14; Fig. 1Aii, mean for population in Fig. 1Aiii), referred to here as “regular tonic.” Second, we observed a single spiking phenotype (Fig. 1Ci) that produced only one spike on the onset despite increasing the amplitude of the current injection (n=16; Fig. 1Cii, population in Fig 1Ciii). These two phenotypes align with those previously described^16,17,32^. In addition, we observed a 3rd phenotype that was characterized by staggered or irregular bursts of action potentials (Fig. 1Bi), which we denote as “chattering” tonic neurons. The bursts increased in spike count and duration with larger current injections (n=14; Fig. 1Bii, population in Fig. 1Biii). The f-I relationship of chattering neurons was more sinusoidal than for regular firing neurons.

**Figure 1.**
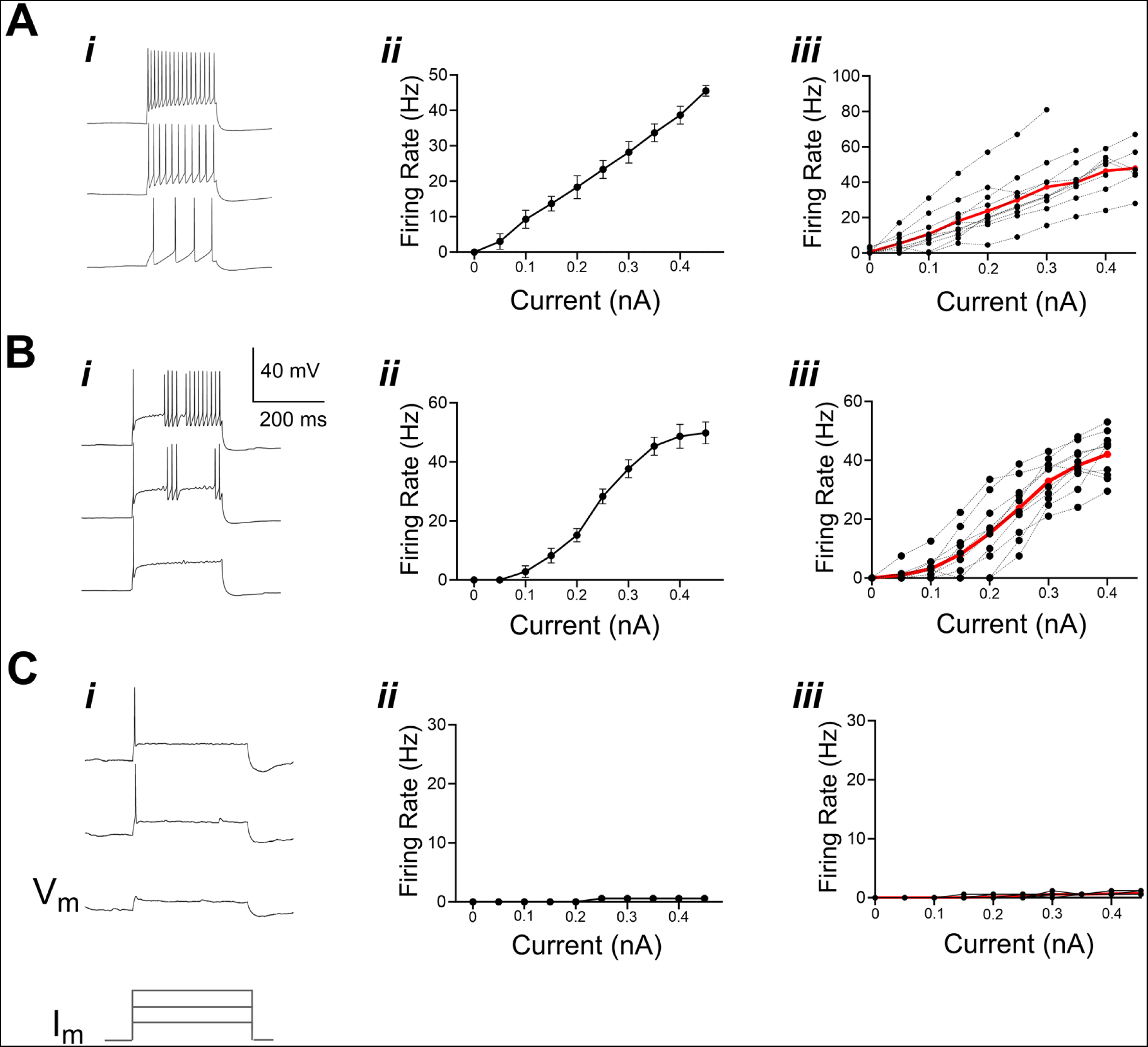
Intrinsic physiological phenotypes of SON neurons. Three phenotypes of responses were identified by their voltage response to current step stimulation: (Ai) regular tonic firing, (Bi) chattering tonic firing and (Ci) single spiking. Qualitative differences were observed in action potential firing patterns (i; traces show voltage responses to current steps of three different amplitudes; bottommost traces in first column illustrate timing and relative amplitude of current steps). Shapes of the firing rate-current amplitude (FI) curves of the two types of tonic firing neurons were linear for regular tonic and sigmoidal for chattering (ii) (mean±s.d. across trials). Matching example individual recordings in columns (i) and (ii); column (iii) overlays FI curves from all individual recordings (black markers) and group averages (red line)(all error bars omitted for clarity). Scale bar A-C: vertical 40 mV; horizontal, 200 ms. Vm, membrane voltage; Im, membrane current.

To determine whether regular firing and chattering neurons could be divided into distinct groups, we quantified the regularity of spiking with the coefficient of variation (CV) (see Methods) of the interspike intervals. The measured CV for the initial step above rheobase appeared to be bimodal with a clear gap at 0.3 (Fig. 2Ai). These groups corresponded to the visually observed chattering and regular phenotypes, respectively, supporting the division of these phenotypes into groups for further analysis.

**Figure 2.**
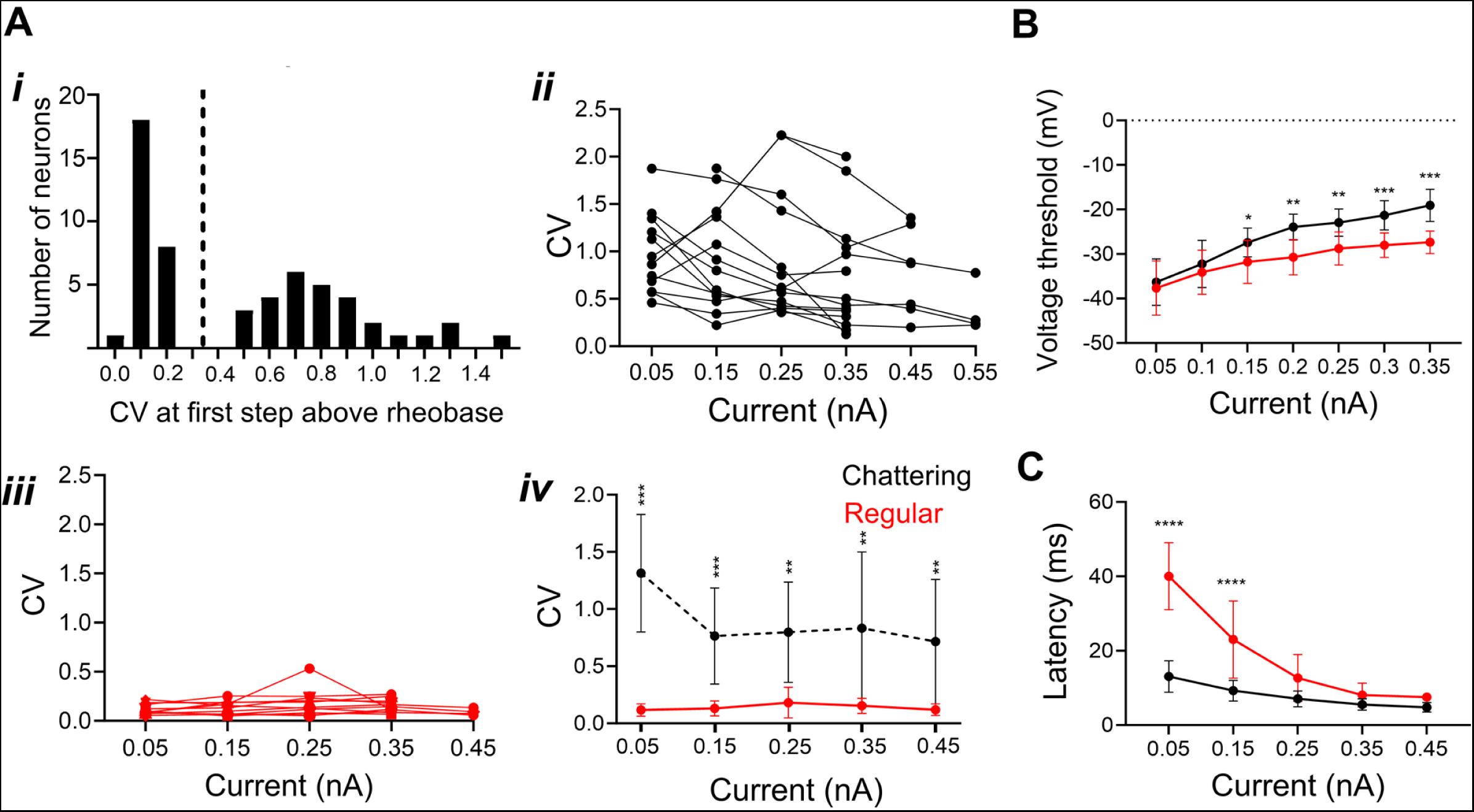
Chattering and regular firing phenotypes were quantifiably differentiable by regularity analysis. A) Histogram of coefficient of variation (CV) of the interspike intervals calculated for the action potentials fired in response to the one current step above rheobase. A clear gap around 0.3-0.4 was used as a criterion for the division of regular and chattering cell types (dashed vertical). CV versus current step amplitude for chattering neurons (Aii; N=14) and regular (Aiii; N=14) neurons. CVs were significantly different between chattering and regular tonic cell types across all stimulus amplitudes, although spiking may regularize with increased stimulus level (Aiv) (2-way ANOVA, main effect by cell type [F (1, 157) = 131.9, P < 0.001]; post-hoc comparisons at each step: P < 0.001 at 0.05 nA; P < 0.001 at 0.15 nA; P = 0.0061 at 0.15 nA; P = 0.0041 at 0.2 nA; P = 0.0029 at 0.25nA; P = 0.0030 at 0.3 nA; P = 0.0070 at 0.35 nA; P=0.0042 at 0.4 nA; P = 0.022 at 0.45 nA, Sidak’s multiple-comparisons test. ***, P<0.001; **, P<0.01. B) Chattering tonic neurons showed an increase in action potential threshold with step amplitude slightly more than regular tonic neurons (ANCOVA, [F[1, 175]=12.31, P=0.004***). C) First spike latency was delayed in regular tonic neurons compared to chattering neurons, which typically fired a spike at the onset. Latency decreased with step amplitude (2-way ANOVA, main effect by cell type [F (4, 84) = 69.37, P < 0.001]; P = 0.001 at 0.05 nA***, P = 0.001 at 0.1 nA***, Sidak’s multiple-comparisons test).

Previous analyses of ‘bursting’ and tonic firing groups have shown that increasing current drive regularizes the firing across the stimulus step^41^. When we analyzed the CV with increasing steps, we found that both the chattering tonic CV (Fig. 2Aii) and the regular tonic (Fig. 2Aiii) did decline, but chattering tonic were still significantly higher than regular firing neurons at all step levels (Fig. 2Ai. 2-way ANOVA, main effect by cell type [F (1, 157) = 131.9, P < 0.001]; P < 0.001 at 0.05 nA, P < 0.001 at 0.15 nA, P = 0.0061 at 0.15 nA, P = 0.0041 at 0.2 nA, P = 0.0029 at 0.25nA, P = 0.003 at 0.3 nA, P = 0.007 at 0.35 nA, P=0.0042 at 0.4 nA, P = 0.022 at 0.45 nA, Sidak’s multiple-comparisons test).

To determine if the onset mechanics of spikes differed between these two phenotypes, we quantified the threshold (Fig. 2B) and the latency to first spike after current injection (Fig. 2C). Interestingly, chattering tonic neurons had comparable thresholds with regular tonic neurons at lower current levels but significantly lower latency. Increasing current increased the threshold in both phenotypes, however the slope of the threshold sensitivity (i.e. the change in threshold with input current) in chattering neurons (slope = 55.7 mV/nA) was significantly higher than the slope for regular tonic neurons (slope = 32.82 mV/nA) (ANCOVA, [F [1,175] = 12.31, P=0.004). Increasing current level also reduced the latency in both phenotypes, with the latency in regular tonic significantly higher at lower current levels than chattering tonic. (2-way ANOVA, main effect by cell type [F (4, 84) = 69.37, P < 0.001]; P = 0.001 at 0.05 nA, P = 0.001 at 0.1 nA, Sidak’s multiple-comparisons test). That difference became non-significant at higher current levels.

We then measured the intrinsic membrane properties of each SON phenotype (Table 1). Chattering neurons had lower whole cell capacitance (n=26, WCC= 50.4± 10.3 pF) than regular tonic (n=33, WCC =69.5±18.8 pF) or single spiking phenotypes (n=23, WCC =57.6±18.6 pF) (Student’s t-test, P=0.0028 chattering vs. regular, P=0.0042, chattering vs single spike, P<0.001, regular vs single spike). Single spiking neurons had the largest afterhyperpolarization (AHP) (12.0±5.3 mV). Chattering tonic neurons had the second largest AHP (4.99±1.19mV) and regular tonic had the smallest amplitude AHP (2.09±2.86 mV) (Student’s t-test, P= 0.0016 chattering vs. regular, P<0.001, chattering vs single spike, P <0.0001, regular vs single spike). Finally, single spiking cells had the highest rheobase (0.30±0.11 nA), chattering tonic had the second largest (0.23±0.12 nA), and regular tonic had the lowest (0.12±0.007 nA) (Student’s t-test, P= 0.003 chattering vs. regular, P<0.001, chattering vs single spike, P <0.0001, regular vs single spike).

**Table 1:** Passive membrane and firing properties.

| Cell Type | Resting Potential (mV) | $R_{in}$<br>(MΩ) | Capacitance<br>(pF) | Rheobase (nA) | Afterhyper-polarization<br>(mV) |
| --- | --- | --- | --- | --- | --- |
| Chattering (n = 26) | -57.8 ± 4.8 | 196.6 ± 26.1 * | 50.4 ± 10.3 * | 0.23 ± 0.12 * | 4.99 ± 1.19 * |
| Regular (n = 33) | -61.9 ± 4.8 | 222.9 ± 28.5 * | 69.5 ± 18.8 * | 0.12 ± 0.07 * | 2.09 ± 2.86 * |
| Single spike (n = 23) | -59.6 ± 4.3 | 147.4 ± 24.3 * | 57.6 ± 18.6 * | 0.30 ± 0.11 * | 12.0 ± 5.3 * |

Mean ± s.d. Input resistance was calculated from the slope of the $I/V$ function in response to hyperpolarizing current injection.
Statistically different from all other types (Student's T-Test, $P < 0.01^{**}$ )

### SON neurons display varied levels of temporal reliability

We next wanted to explore if intrinsic membrane properties of the three SON phenotypes engendered different levels of temporal capabilities. To investigate how well the SON neurons locked to the current stimulus, we pharmacologically blocked synaptic activity (see *Materials & Methods*) and measured their responses to frozen noise over repeated trials to analyze the spike timing. We recorded timing data from 78 SON neurons: 33 regular tonic, 20 single spiking, and 25 chattering tonic neurons. A single DC current step amplitude was chosen to generate firing levels in the tens of Hz range. White noise was added to the current step amplitude (4σ, see Methods). All SON phenotypes produced reliable firing patterns upon noisy current injection (Fig 3Ai-iii, iv). The trial-to-trial temporal firing reliability was analyzed using a shuffled autocorrelogram (SAC), a histogram of all across-trial interspike intervals, excluding within-trial spike pairs (Fig. 3Bi-iii). An example voltage trace is shown in figure 3C. The peak value (measured as correlation index, or CI) and width of the SAC indicate the degree and precision of the time locking of the spikes across trials, respectively. Single spiking neurons were the most reliable, with the highest CI (n=20, mean = 81.34±24.9) and the narrowest SAC (FWHM mean = 1.04±0.53 ms). Chattering tonic neurons displayed the second highest CI values and the second narrowest SAC (n=25, CI mean = 26.11±9.17, FWHM mean = 2.7±1.11 ms). Regular tonic neurons had the lowest CI values and the widest SAC (n=33, CI mean = 7.33±2.58, FWHM = 7.53±1.99 ms). Both the CI (one-way ANOVA-[F (2, 59) = 109.7], P<0.001) and the FWHM (one-way ANOVA [F (DFn, DFd) = 8.731 (2, 59)], P<0.001) were statistically different between the three phenotypes (Figure 3Di, ii). These results suggest that different SON phenotypes are intrinsically capable of encoding differing levels of temporal information.

**Figure 3.**
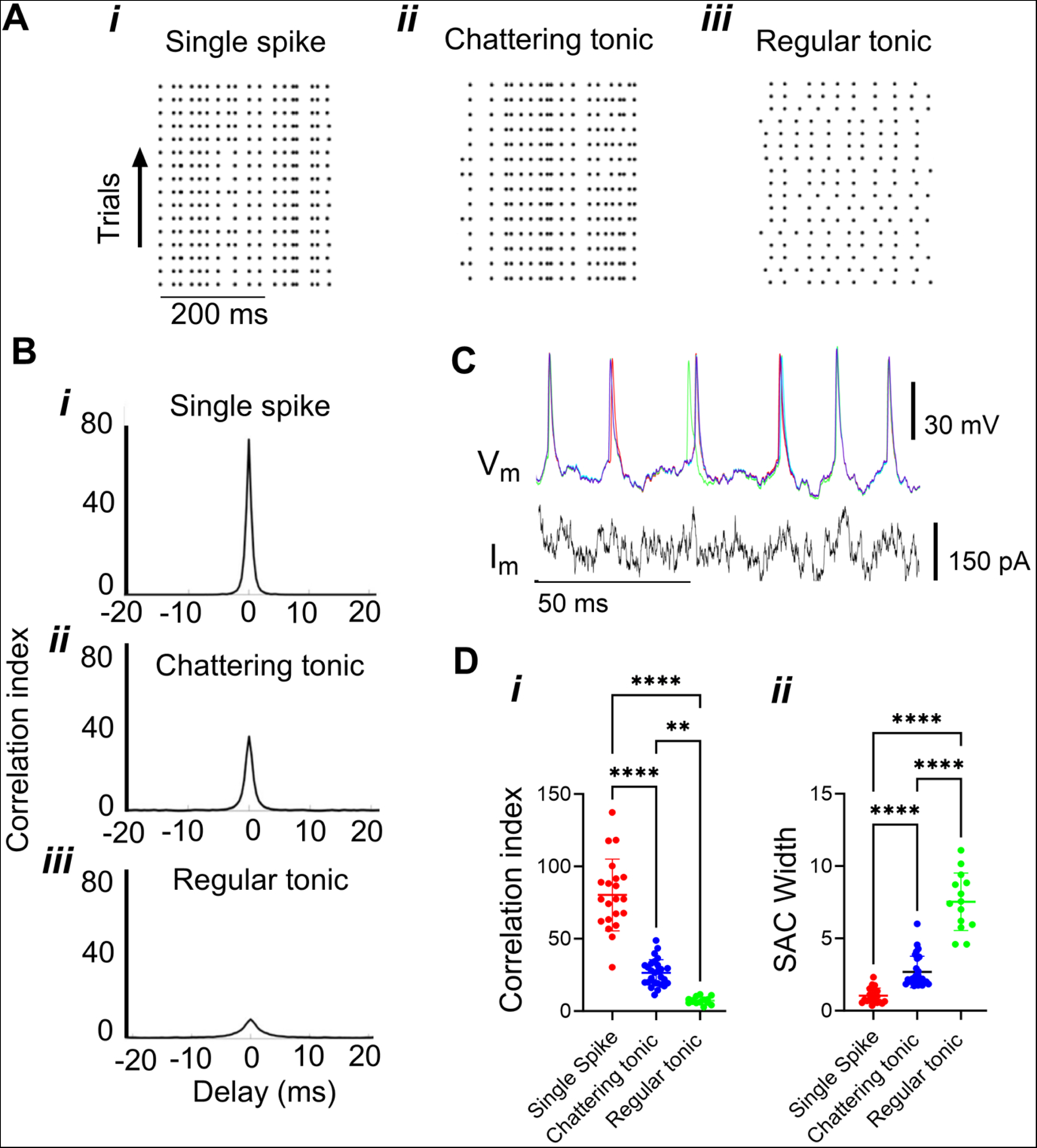
Spike timing reliability depends on cell type. A) Raster plots in response to a noisy current stimuli (not shown). An example neuron (i-iii) of each cell type show reliable firing to repeated presentations of the same stimulus B) Example of a shuffled autocorrelogram (SAC) for each cell demonstrated in A. C) Voltage responses (*Vm*) of repeated iterations of identical noisy current injections (*Im*) from a chattering tonic neuron. D) Quantifying the SAC parameters, peak correlation index (CI) and full-width at half-max (FWHM) of each cell type. *i* Cell types in SON have distinct CI peaks: *i* single spiking cells, CI=81.34±24.9, n=20; *ii* chattering tonic, CI= 26.11±9.2, n=25; *iii* regular tonic neurons CI=7.33±2.7, n=33. One-way ANOVA [F (2, 59) = 109.7], P<0.001***). *ii* Cell types in SON have distinct autocorrelogram (single spiking cells, FWHM = 1.04±0.53 ms; chattering tonic FWHM = 2.7±1.11 ms; regular tonic cells, FWHM = 7.53±1.99ms, (one-way ANOVA [F (2, 59) = 123.6, P<0.001***)

### SON tonic phenotypes exist along a spectrum of fluctuation sensitivity

We next considered the effects of naturalistic noisy current injections on the input-output functions of the two types of tonically firing SON neurons. Characterizing noise sensitivity means quantifying neurons on a spectrum of integrators to differentiators. Fluctuation insensitive neurons are integrators, which increase spiking monotonically with current but not with noise level. Comparatively, fluctuation sensitive neurons behave are differentiators, increasing spiking with current level, but also with noise level^39,41,43^. We recorded 69 neurons for these experiments: 30 regular tonic neurons and 39 chattering tonic neurons. The addition of noise altered the input-output functions of chattering tonic neurons (Fig. 4A), with enhanced fired to the larger voltage swings in the noisy currents, thus responding as differentiators. The addition of noise did not later the input-output functions of the regular tonic neurons, whose firing rates were indifferent to the noise level, thus responding as integrators (Fig. 4B). To quantify the noise responses of tonic SON neurons, we measured the area of the curve between the highest and lowest levels of noise at each current step (Fig. 4C*i*) and as well as the difference in firing between high and low noise levels at the highest level of current. We determined that chattering and regular tonic neurons displayed different levels of fluctuation sensitivities (Fig. 4Cii, chattering tonic ADI mean = 0.35, n= 39, regular tonic ADI mean= 0.08, n =30, unpaired t-test, P<0.001). These results suggest that a subset of tonic neurons in SON preferentially encode temporal variations in their input, while others remain invariant.

**Figure 4.**
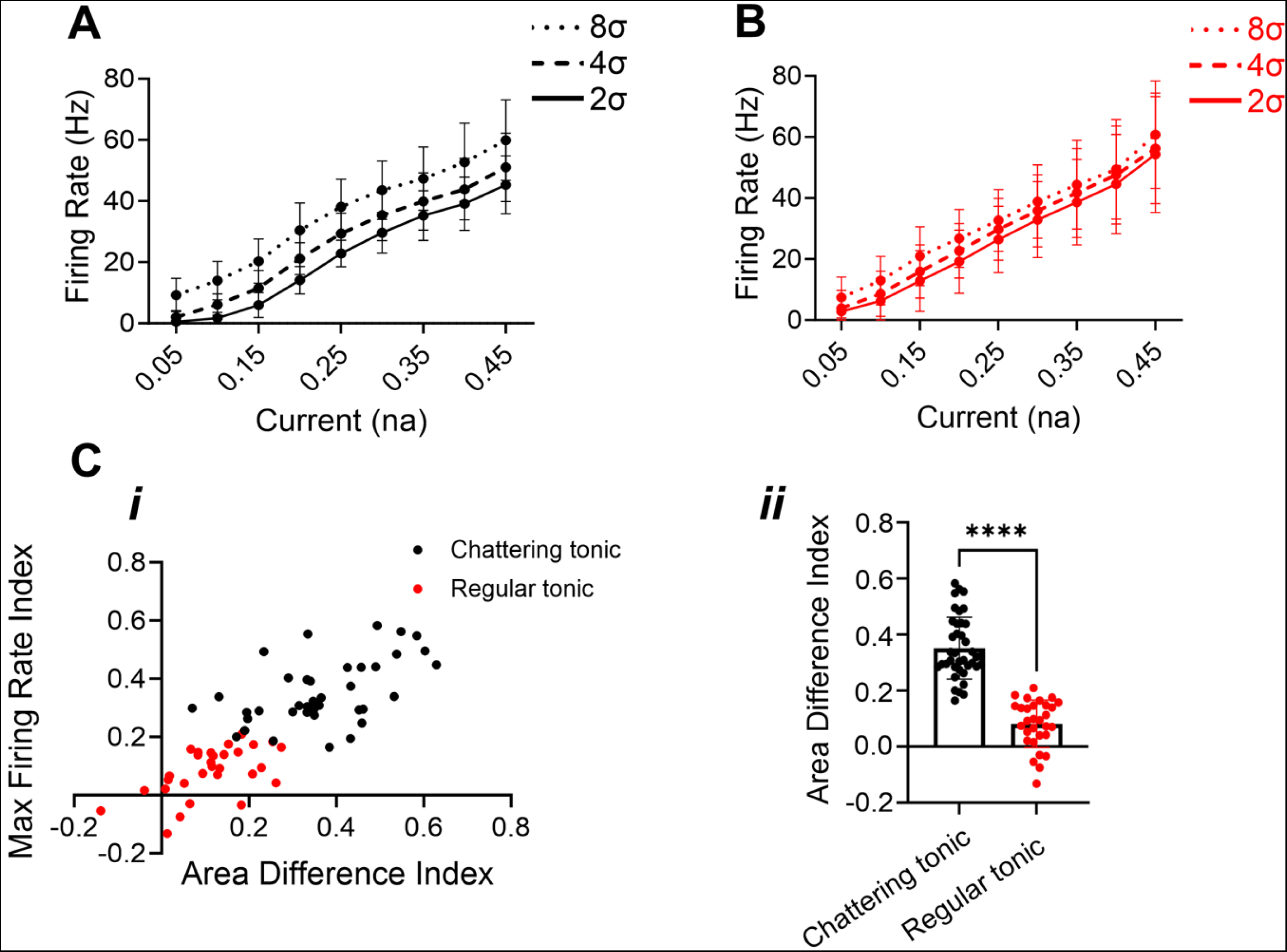
Temporal fluctuation sensitivity depends on cell type. FI curves of tonic firing SON neurons in response to noisy currents were measured at three different noise fluctuation amplitudes (standard deviation around the mean: 2σ,4σ or 8σ; see Methods) were measured. Abscissa represents the mean current amplitude for each stimulus. (A) In one chattering neuron, increasing the noise fluctuation amplitude elicited higher firing rates. (B) In one regular neuron, there was little effect of noise amplitude on the firing rates (B). Ci) Comparative analysis of FI curves sensitivity by plotting the difference in maximum firing rate and the integrated difference between the curves for largest (8σ) versus smallest (2σ) noise conditions (see Methods). Chattering and regular tonic neurons comprised different ends of a continuum of sensitivity to fluctuations. Cii) Grouped values were statistically different (chattering, N=39; regular, N=30, Student’s t-test, p<0.001).

### SON neurons exited nucleus in at least three distinct fiber tracts and correlated with physiological phenotype

Here, we used anatomy to explore the anatomy of the fiber tracts that exited SON neurons. We recorded from individual neurons *in vitro* and dialyzed cells with biocytin. We visualized the biocytin and were able to reconstruct the neurons, with a focus on reconstructing the cell body and axon. We reconstructed neurons using *Neurolucida* software and were able to identify axons from their uniform thickness and lack of branching. Axons needed to extend a minimum of 400 µm beyond the boundary of the SON to be included in the analysis. Thirty-five neurons were recovered whose reconstructed axonal projections met the criteria and for which physiological data were collected: 14 single spiking, 10 regular tonic, and 11 chattering tonic neurons.

To visualize populations or clusters, all reconstructed neurons were superimposed onto a single representative left SON (Fig. 5Ai). Three fiber tracts were visually distinguishable. We quantified these projections by measuring the angle between the center of the cell body with the point on the SON boundary where the axon crossed, with due lateral as a reference angle of 0° (Fig.5Aii) Three distinct fiber tracts could be differentiated on the basis of this crossing angle: an ipsilateral lateral tract (between 30° and 70°, n=5), an ipsilateral medial tract (between 90° and 135°, n=17) and a contralateral tract (between 180° and 280°, n=13).

**Figure 5.**
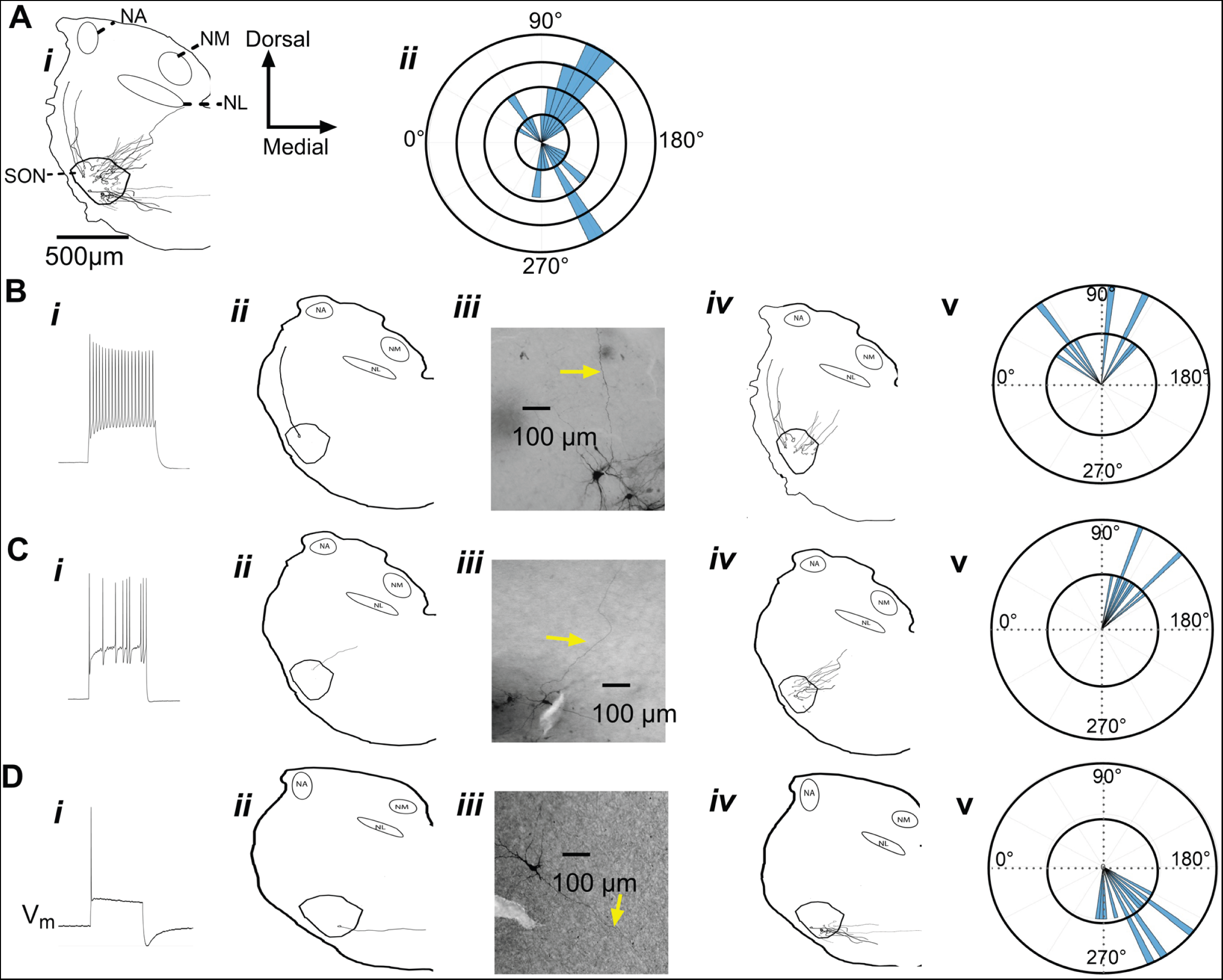
Morphological analysis of axonal projection patterns. Physiologically recorded SON neurons were intracellularly labeled with biocytin, processed and reconstructed partial morphology to locate the cell body and determine the projection pattern of the axon as it leaves the SON. Axons appeared to exit the SON in three directed fascicles. A) (Top Left) Compiled overlay of 39 separate recovered neuron in 30 slices onto a typical anatomical outline of the ventral left quadrant of the chick brainstem. B) (Top Right) When the angle of the efferent axon was measured to quantify these tracts such that lateral = 0°, dorsal = 90°, we could identify an ipsilateral lateral tract (30° ≤ x ≤ 70°; n=5), an ipsilateral medial tract (90° ≤ x ≤ 135°; n=17) and a putative contralateral tract (between 180° ≤ x ≤ 280°; n=13). Radius indicates axon counts (each concentric ring = 1 count). C) Physiological cell types showed preferential efferent projection angles. For each cell type, example voltage traces (i), reconstructed axon shown relative to SON and brain stem outlines (ii), photomicrograph of the cell body and axon (iii), polar plot of axonal projections of cell type group (iv), and overlay summary of all axon reconstructions for the group (v) polar histogram quantifying the distribution of axons exiting the SON at particular angles. D) Axons from regular tonic cell type fell within one of the two ipsilateral tracts. E) Axons from chattering tonic cell type SON neurons all fell within the ipsilateral medial tract. F) Axons from single spiking SON neurons all fell within the contralateral tract.

To explore the relationship between SON response types and postsynaptic targets, we matched 35 reconstructed neurons with the data from electrophysiological recordings and found that specific SON phenotypes underlie the separate observed fiber tracts. Chattering tonic neurons exited solely via the ipsilateral medial projection (n=11) (Fig. 5A*i-iv*). Single spiking neurons comprised the entirety of the contralateral projection (n=14) (Fig. 5B*i*-*iv*). Finally, regular tonic neurons comprised the entirety of the ipsilateral lateral projections (n=5), however, they also exited via the ipsilateral medial projections (n=5) (Fig. 5C*i-iv*). These results suggest that SON phenotypes underlie the divergent projections.

### Cells in SON are clustered by their electrophysiological phenotype

We then asked if SON phenotypes were organized within the anatomy of the nucleus. We analyzed the fills of neurons that met the axon length criteria for the projection analysis (Fig. 6- *top row.* n=35) and the cell bodies of all filled neurons (Fig. 6- *bottom row*. n=135). These clusters revealed some loose organization of SON phenotypes. Single spiking neurons tended to exist in the ventral areas of the SON (12/14 of neurons that had sufficient axon length Fig. 6A*i*, 33/45 of all single spiking neurons), while both tonic phenotypes tended to exist in the dorsal area. For chattering tonic neurons, 30/46 existed in dorsal areas. Regular tonic neurons ended up being identical in proportion, with 30/46 existing in dorsal sections of the SON. These results indicate that SON neurons tend to cluster within regions of the SON.

**Figure 6.**
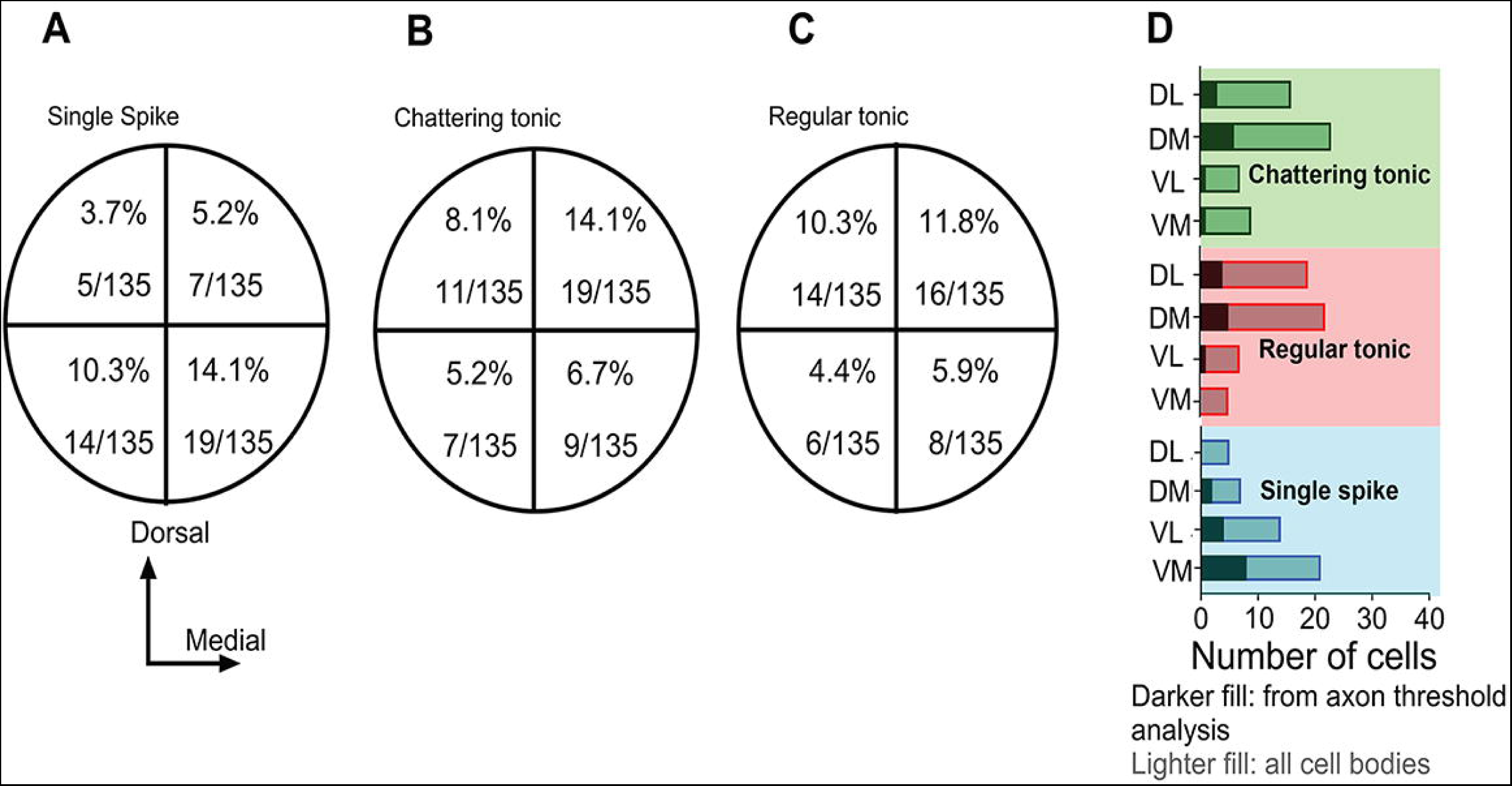
Distribution of the location of cell bodies within SON differs with physiological type. A-C) Diagram of the coronal section of SON into dorsoventral and mediolateral quadrants with the number of cell bodies located in each out of a total of 135 recovered and corresponding physiological types. Percentages expressed as percent of total. D) M, Medial; L, Lateral; D, Dorsal; V, Ventral. Same data as a bar plot shows the relative distributions of both tonic types were similar and preferentially in dorsal quadrants, while single spiking neurons were preferential in ventromedial. Distribution of SON neurons for which axonal reconstructions were recovered as well and shown as darker shaded regions within bars and had a similar distribution.

### Rostrocaudal distribution of SON phenotypes

The three observed fiber tracts exited SON along a rostral-to-caudal axis. Using cytoarchitectural references (NL, NM, NA, and medial vestibular size), we split these fiber tracts into caudal, medial, and rostral sections. 11/17 (64.7%) of ipsilateral medial axons and a 1/ 5 (20%) of the ipsilateral lateral axons exited in the most caudal plane. 5/17 (29.4%) of the ipsilateral medial axons, 4/ 5 (80%) of ipsilateral lateral axons, and 1/16 (6.3%) of contralateral axons exited in the medial plane. 1/17 of the ipsilateral medial axons and 15/16 contralateral axons exited in the rostral plane (Fig. 7C). These results suggest that the axons of SON neurons are organized and “bundle” into specific planes, depending on their postsynaptic targets.

**Figure 7.**
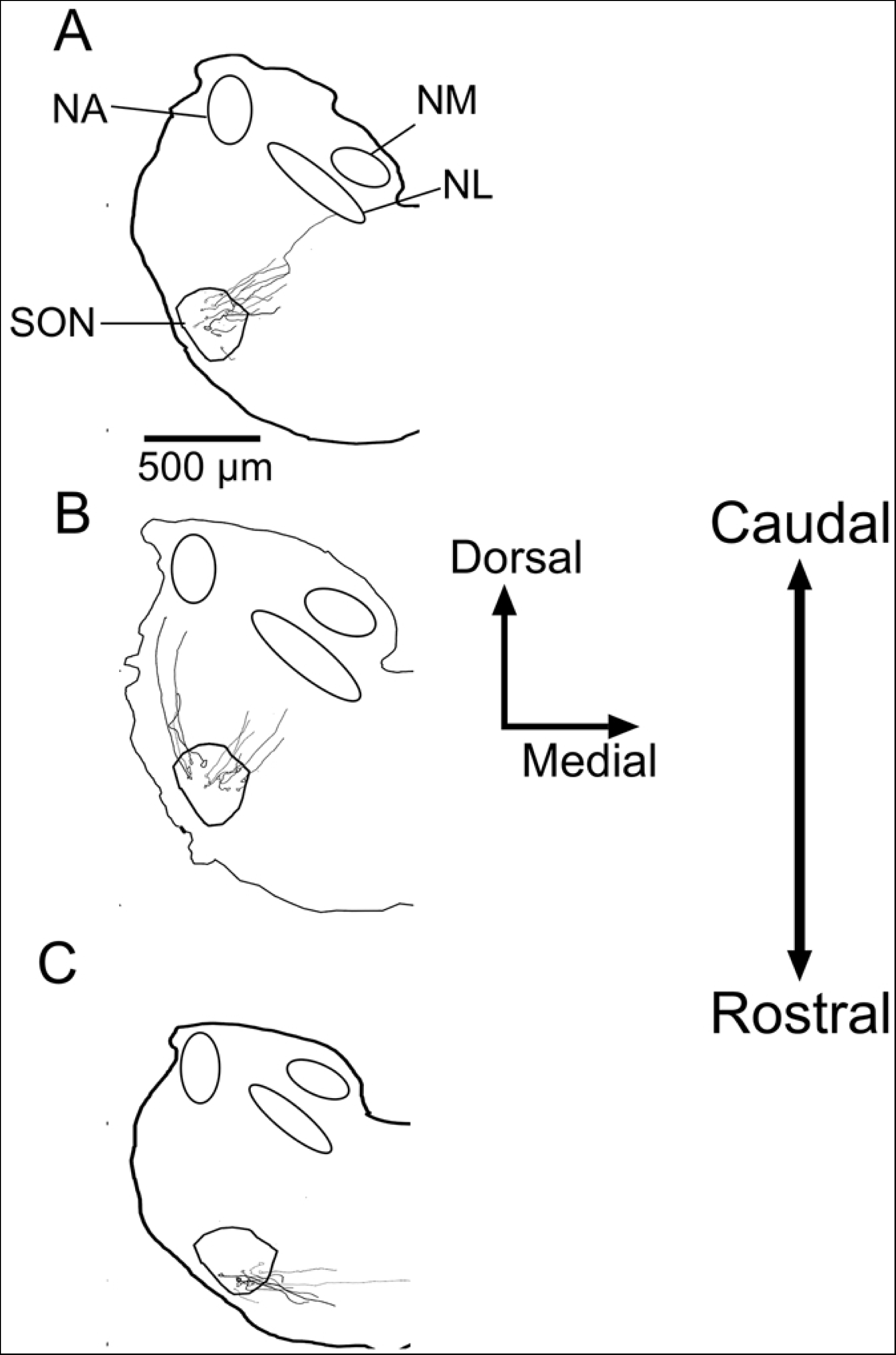
Axonal projection patterns differed across the rostrocaudal extent of SON. Nuclei are labeled. A) In the caudal most sections axonal projections were mostly via the ipsilateral medial tract (10/10) ipsilateral medial, 4 regular tonic & 6 chattering tonic) B) Middle sections contained axonal projections either via the ipsilateral medial tract (8/13, 5 chattering tonic and 3 regular tonic) or the ipsilateral lateral tract (5/13, all 5 regular tonic) C) Rostral most sections contained axonal projections almost entirely oriented contralaterally (16, all single spiking). These patterns are consistent with relative rostrocaudal location of NM and NL (more caudally), NA (more rostral) versus the SON (most rostral). N.b. while SON is located more rostrally than the other brain stem nuclei, sections were cut slightly off coronal (dorsocaudally to ventrorostrally) to maximize the recovery of axonal projections

## Discussion

Neural inhibition contributes to numerous computational functions in auditory brainstem circuits, such as intensity-dependent gain-control^3,10,20^, phase locking^5–10^, ITD, gap detection^11,12^, and amplitude modulation coding^3,13^. Synaptic inhibition in the avian auditory brainstem originates largely from the SON, which projects to the ipsilateral NA, NM, and NL, or to the SONc^31^. These projections arise from distinct SON populations. The inhibition these SON neurons provide is diversified enough to perform all these functions, suggesting underlying specializations within SON. Two phenotypes based on intrinsic physiological firing elicited *in vitro* have been described in the SON: a tonic firing phenotype and a single spiking phenotype^16,17,32^, however the functional role for each phenotype is not clear. We investigated whether specific cell types underlie the divergent projections from SON as a first step toward relating intrinsic properties to circuit function.

### Two tonic firing phenotypes in SON

In this study, we describe a third, previously undescribed, phenotype which we term a “chattering” tonic neuron, distinct from the “regular” tonic neurons. To distinguish tonic firing phenotypes, we quantified each neuron’s interspike interval CV as a measure of the regularity of firing and found a bimodal distribution in the population of tonic neurons that corresponded to the qualitative spiking pattern of chattering and regular. Using CV as a grouping criterion, we tested other electrophysiological properties and found differences in spike latency, rheobase, input resistance, and afterhyperpolarization, indicating that there are two distinct phenotypes within our sample of tonic neurons.

Variations in excitability and spiking properties may emerge from the differential expression, kinetics, and densities of ion channels (See^44^ for summary.) The Kv1 family of channels is a class of low threshold activated voltage-dependent potassium channels that are prevalent in auditory regions, particularly in temporally sensitive areas^45–49^ and are critical for coincidence detection neurons across numerous species^50–55^. While most prominent in avian NL neurons (see review^56^), they are also present in subsets of NA neurons, including distinct tonic phenotypes in which dendrotoxin, a Kv1 channel antagonist, abolished all firing response patterning^39^. The chattering phenotype could also result from sodium channels inactivating after a burst of firing, while Kv1 channel activity acts to prevent further spiking until the sodium channels de-inactivate.

Alternatively, another subthreshold voltage-dependent delayed rectifier potassium current, labeled M-type, contributes to burst firing *in vivo* in the substantia nigra and chattering phenotypes in entorhinal cortex^57,58^, but this current type has not yet been identified as present in avian auditory neurons.

The novel observation of a chattering phenotype in this study compared to previous *in vitro* studies of SON could be due to differences in temperature, age, or stimulus protocol. At near physiological temperatures (or as in our experiments ∼34℃) stimulation can elicit more complex firing behavior compared to that at room temperature^16,32^. Low temperatures slow the kinetics of channels that contribute to variation in spike patterns^59,60^. While age-dependent effects on spiking has also been well-documented^52,61^, this is unlikely to explain the differences here as age ranges in previous studies overlapped with the ages we used here^16,62,63,17,32^. Finally, differences in stimulus protocol could obscure spiking patterns that are most apparent at current levels just above rheobase, as we find with the chattering phenotype. Increasing current leads to an increase in the number of spikes in each burst, eventually leading to tonic firing throughout the duration of the current injection. Yang et al.^16^ did not report their current step intervals, however Carroll & Hyson^32^ reported using intervals smaller or equal to the intervals we used, thus we believe that temperature was a primary driver in the differences in phenotypes observed.

### SON neurons encode temporal features of sound

The capability of neurons to encode temporal features is dependent on synaptic dynamics^62–64^ and intrinsic membrane properties^41,42,65,66^. Temporal features emerge at different timescales. Phase locking, or the firing at a particular phase of a modulated stimulus, is important for pitch perception^67^ and for binaural ITD computations^68^.

Envelope coding, or firing to instantaneous changes in amplitude, is important for sound recognition^69–71^. Historically, SON neurons were not thought to encode temporal features and merely rate code, but a subset of *in vivo* recordings from SON neurons showed phase locking in neurons up to 4 kHz. However, the strength of the phase locking varied considerably amongst neurons^17^.

Our study investigated intrinsic sensitivity of SON neurons to temporally modulated inputs *in vitro* using fluctuating noisy current injections^72^, where fluctuation size correlates with synchronous synaptic release. Each SON phenotype was found to have a different level of temporal sensitivity, with single spiking neurons being the most reliable, regular tonic neurons the least reliable, and chattering tonic neurons in between. These different phenotypes may underlie the spectrum of temporal sensitivity demonstrated in Coleman et al.^17^. We also varied the level of fluctuation to determine if SON neurons differentially responded to fluctuation size. We found that chattering tonic neurons tonic neurons increased their firing with larger fluctuations but regular tonic neurons did not. These results suggest that input synchrony, as well as mean input, drive the level of inhibitory output from the SON.

### SON neurons provide inhibition to divergent postsynaptic targets organized by intrinsic spiking properties

In previous studies, retrograde labeling revealed distinct SON populations that send axons either ipsilaterally towards the lower order nuclei (NA, NL, and NM) or to SONc^31^. Whether these divergent projections represented different electrophysiological types was not known. To address this issue, we reconstructed the axons of recorded neurons and found a clear pattern. Each neuron had an axon that exited the SON in one of three directions: as an ipsilateral projection that followed the olivary-trapezoid tract toward NM and NL, an ipsilateral projection that followed a lateral tract, potentially in the same region as the inferior cerebellar peduncle toward NA, or a contralateral projection fiber tract, which crosses along the trapezoid body decussation that headed toward SONc^73^. The firing response phenotype of each neuron correlated with its preferred axonal fiber projection: axons in the ipsilateral medial projection originated from both chattering tonic and regular tonic neurons; axons in the ipsilateral lateral tract originated exclusively from regular tonic neurons; and axons in the contralateral projection originated exclusively from single spiking neurons.

### Role of inhibition in the avian auditory brain stem

The critical role of inhibition in sound encoding by the timing nuclei has been well established. Blocking inhibition worsens phase locking in NM^7,74^ and compresses the dynamic range, or range of sound intensities encoded before saturating, of low frequency NM neurons^10^. In NL, inhibition shortens the window for coincidence detection, improving ITD calculation^16^. Nishino et al.^20^ demonstrated that low frequency NL neurons (<1 kHz) had improved contrast between their best and worst frequencies during increased sound levels due to increased levels of inhibition. Inhibitory postsynaptic currents (IPSCs) in NM are slow and temporally summating^5,16,75–77^, however the kinetics of IPSCs are more rapid in NL than NM, indicating a preference towards maintaining temporal structure^75^. Although summation effects would blunt temporal precision, the arrival of SON-driven synaptic inputs from temporally-sensitive chattering tonic neurons could influence temporal processing in NL, especially in low frequency regions.

Less is known about the function of inhibition in the so-called intensity pathways of avian brainstem, but some evidence from synaptic studies suggests differential processing. Kuo et al.^75^ demonstrated the kinetics of IPSCs in NA (putative SON inputs) are more rapid than in either NM or NL and vary amongst tonic and single spiking neurons in NA. The function of inhibition in NA has not yet been studied using pharmacological methods, but feedback integrator-like regular tonic SON neurons suggests a negative gain-control as a primary function. Such a gain control function would be consistent with the projection of the regular tonic, ‘integrator”-like neurons found in our study. On the other hand, the rapid synaptic kinetics could indicate a role in spectrotemporal processing which would be consistent with the enhanced envelope coding in NA compared to their nerve inputs or that in the timing^40,42^

Coleman et al.^17^ demonstrated both glycinergic and GABAergic signaling between the two SON and that pharmacologically blocking either increased the firing rate to sound stimuli, while blocking glycinergic signaling worsened phase locking. Binaural coupling of the two SON is theorized to be critical for ITD processing within NL, particularly during large interaural level differences (ILD) (see review by Burger et al.^78^). In short, NL must differentiate excitatory drive from a strong monaural stimulus or a well-timed binaural stimulus. Contralateral auditory drive disinhibits NL, boosting ITD computation. Our study suggests that the primary source of contralateral inhibition may be the single spiking neurons from the opposite SON. The phasic and temporally locked nature of these single spiking neurons could selectively bolster activity being used for ITD detection during large contrasts of ILDs.

Important to note, the location of cell bodies of these neurons within the SON lined up with previous retrograde labeling experiments^31^, with single spiking cells positioned ventrally and tonic neurons positioned dorsally.

A limitation of this experiment lies in the inability to follow axons entirely to their postsynaptic target. It is possible that axons could turn and synapse in a location that is not apparent to by the length of axon measured. Burger et al.^31^ demonstrated that ipsilateral projections from SON provided input to all three of the lower brain stem nuclei, NA, NL, and NM. Therefore, this ipsilateral lateral projection could synapse on NM and NL as well, though how that is accomplished is not clear. Alternatively, this population could project outside of the brainstem.

### Mammalian vs avian inhibitory architecture

Birds utilize sounds to accomplish the same sensory tasks as mammals, but with simpler circuitry^79^. In birds, inhibition in the auditory brainstem primarily originates from the SON. Importantly, there is no local inhibition. Comparatively, inhibition in the mammalian auditory brainstem arises from multiple sources and is utilized in several different processes^9,18,80–85^. D-stellate cells in the ventral cochlear nucleus (VCN) utilize large dendritic arbors to span auditory nerve fibers and provide broadband local inhibition, the dorsal cochlear nucleus (DCN), and to bushy cells in the contralateral VCN^86–89^. Tuberculo-ventral cells in DCN receive AN input and project to the ipsilateral VCN^90^. Finally, L-stellate cells in the VCN receive narrowband inputs and project to ipsilateral T-stellate and bushy cells^18^.

A functional difference in the use of inhibition is in ITD calculation. Mammalian ITD circuits are thought to have evolved after ILD circuits, due to an increase in body and head size^79^. In mammalian ITD processing, excitatory inputs from bushy cells in the VCN drive activity in contralaterally projecting glycinergic neurons, shifting the ITD preference towards the contralateral ear^91,92^. Avian ITD circuits strictly use excitatory feed-forward circuits for coincidence detection while inhibition modulates the process. Additionally, interaural level difference (ILD) calculations occur outside of the brainstem within birds, occurring within lemniscal nuclei^93–95^, while mammalian ILDs are calculated largely in the brainstem^96^.

## Grants

This work was supported by the NIH Grant R01DC1000 (KMM) and by T32 DC-00046 (JFB) from the National Institute of Deafness and Communicative Disorders of the National Institutes of Health.

